# Effects of land-use intensification on nectar resources, bird assemblages and bird-flower interaction networks in the Western Ghats, India

**DOI:** 10.1101/2025.10.01.679556

**Authors:** Nandita Madhu, Vishal Sadekar, Rajah Jayapal, Rohit Naniwadekar

## Abstract

Agricultural expansion is a major driver of tropical forest loss globally, making it critical to determine factors that maintain key plant-animal interactions, such as nectarivory, in human-dominated landscapes. In the Paleotropics, which is experiencing rapid expansion of tree crop plantations, few studies have compared floral and nectar-feeding bird communities, their interactions, and network organisation along a tree crop intensification gradient. In a cashew agriculture-forest matrix of the northern Western Ghats of India, we compared forests, cashew agroforests, and cashew monocultures to assess, 1) the abundance, diversity, and composition of flowers and nectar-feeding birds, 2) diversity of bird-flower interactions, and 3) network organisation. Floral abundance declined most sharply in monocultures, followed by agroforests. The abundance and diversity of nectar-feeding birds peaked in agroforests. Forests had the highest floral diversity and bird-flower interactions. Nectivore community composition differed across land-use types, with agroforests harbouring both forest specialists and open-habitat species typically found in monocultures. We found that sunbirds dominated plant-flower interactions across land-use types. After rarefaction (to control for differences in network size), agroforests and monocultures showed higher connectance, specialisation, and modularity than the forest network. Our results suggest that plantations that retain native trees (agroforests) can partially offset the alterations caused in floral resource availability and nectar-feeding bird communities by cashew cultivation. Planting bird-pollinated native tree species along orchard edges can make cashew farming more suitable for nectar-feeding birds, and has the potential to benefit cashew production by attracting avian pollinators.

## 2. INTRODUCTION

Tropical forests constitute almost half of the world’s forests (FAO, 2020) and harbour more than 50% of the world’s known species, including 90% of terrestrial bird species (Barlow et al., 2018; Shennan-Farpón et al., 2021). However, they face multiple threats, with deforestation, particularly for conversion to agriculture (Pendrill et al., 2022), being the primary driver of biodiversity loss (Maxwell et al., 2016; Morris, 2010). Given large-scale agricultural intensification, it becomes increasingly important to assess biodiversity responses to habitat modification and determine factors that may enable biodiversity persistence in human-dominated landscapes. While open agricultural systems are detrimental to forest biodiversity, tree-based agriculture is generally considered less harmful (Felton et al., 2010; Wang et al., 2022). While the impacts of monoculture plantations, which seldom retain native tree cover, on birds have been assessed in tree crops such as oil palm and rubber (Azhar et al., 2014; Tejeda-Cruz & Sutherland, 2004; Yahya et al., 2017), there has been an increased interest in determining the conservation potential of “wildlife-friendly” agroforestry plantations that retain some native tree cover (Bhagwat et al., 2008). Agroforests have been reported to reduce biodiversity losses compared to monoculture tree plantations (Alvarez-Alvarez et al., 2022; Warren-Thomas et al., 2020). While bird community responses to agroforests and monoculture plantations have been well documented (Birch et al., 2024; Raman et al., 2021), responses of specific dietary guilds, such as nectar-feeding birds, remain relatively less explored.

Nectar-feeding birds are the most speciose of vertebrate pollinators in the world (Cronk & Ojeda, 2008; Fleming & Muchhala, 2008). However, bird pollinators have faced substantial increases in extinction risk in recent decades (Regan et al., 2015). Given their contribution to supporting local food security through the pollination of agricultural crops (Requier et al., 2023), it is important to assess the key determinants that enable nectarivores to persist in human-modified systems thereby contributing to agricultural sustainability and conservation efforts globally.

While nectar-feeding birds are generally negatively influenced by anthropogenic disturbances (Gray et al., 2007), some evidence suggests that their abundance and richness may be higher in agroforests than in other systems (Jarrett et al., 2021). This has been attributed to the greater availability of nectar resources in agroforestry systems (Udawatta et al., 2019) and the presence of native tree cover (Alvarez-Alvarez et al., 2022). Nectar-feeding birds often track floral resources spatially and temporally (Bennett et al., 2014; Cotton, 2007; Pollock et al., 2020). Monocultures may provide abundant nectar during the crop flowering, provided that its flowers are visited by birds. This contrasts with forests, which can offer diverse resources to nectar-feeding birds over extended periods. Assessing resource availability and diversity across increasing levels of tree crop intensification and linking them with nectar-feeding bird resources and determining its implications for bird-flower communities can offer valuable insights for the management of agroforests and tree-crop monocultures.

Conversion of forests to tree-crop plantations is associated with the loss of forest specialists and the influx of open-habitat species, driven by environmental filtering. This often results in compositional turnover, with generalists replacing specialists (Castaño-Villa et al., 2019; Madhu et al., 2025). In contrast, agroforests characterised by intermediate disturbance, can support higher diversity as they harbour birds from forests and open habitats (Edo et al., 2024). Such plantations are structurally and functionally complex, with the presence of an understory as well as canopy trees (Madhu et al., 2025). This loss and gain of species is better assessed by examining compositional changes in flowers and nectar-feeding birds, complementing richness and abundance comparisons across land-use types.

Interactions between nectar-feeding birds and flowers are influenced by neutral processes, such as abundance of nectar resources (Schmid et al., 2016), and niche-based processes, like flower morphology and competition (Rico-Guevara et al., 2021; Uceda-Gómez et al., 2024). Alterations in abundance and composition of flower resources and nectar-feeding birds due to habitat conversion can have cascading impacts on community organisation. Community organisation can be best studied through examining properties of bipartite networks. The re-wiring of interactions in modified habitats can affect network metrics like connectance, specialisation, nestedness, and modularity. While the impacts of agriculture on interaction networks have been mostly assessed for invertebrate pollinators (Adedoja & Kehinde, 2018; Shinohara et al., 2019), vertebrate-mediated pollination, particularly those involving nectarivorous birds, remain comparatively less studied (but see López-Flores et al., 2024; Morrison & Mendenhall, 2020).

Studies assessing responses of nectar-feeding birds to forest conversion, fragmentation, and loss, have mostly been conducted in the Neotropics with a primary focus on hummingbirds (Bustamante-Castillo et al., 2020; López-Flores et al., 2024). Insights on sunbirds, their counterparts from Africa, Asia, and Australia, are relatively scarce (but see Mgimwa et al., 2025). Although not as morphologically specialised as hummingbirds (Fleming & Muchhala, 2008; Stiles, 1981), sunbirds display considerable variability in morphological and behavioural characteristics, allowing them to feed on diverse array of floral and invertivore resources (Janecek et al., 2021). They constitute the major group of avian pollinators (along with spiderhunters) in Asia, Australasia, and Africa and have been observed pollinating crops such as tea (Sun et al., 2017) and banana (Itino et al., 1991). However, assessing the responses of the overall nectar-feeding bird community, that includes groups such as flowerpeckers, leafbirds, and prinias (Ali and Ripley, 1999), can provide a comprehensive understanding of how bird-mediated nectarivory is influenced by land-use modification.

In the northern portion of Western Ghats - Sri Lanka Biodiversity Hotspot, < 5% of the land is under the Protected Area network with most forests being privately owned. These forests are being converted to cashew monoculture plantations at an alarming rate (Chhaya et al., 2025; Rege et al., 2022) making it critical to determine factors that may enable biodiversity to persist in human-dominated landscapes. Across a cashew-intensification gradient of the northern Western Ghats of India, we compared across forests, cashew agroforests, and cashew monocultures, 1) abundance of flowers over and nectar-feeding birds, 2) diversity of flowers, nectar-feeding birds, and nectarivorous interactions, 3) composition of flowers and nectar-feeding birds, and 4) plant-nectar-feeding bird network organisation. By integrating data on resource availability, nectar-feeding bird abundance, diversity, composition, interactions, and network organisation across land-use types, we aim to better understand the effects of land-use change on bird communities and ecosystem functioning.

## 3. METHODS

### 3.1. Study Area

We conducted this study between January and April 2004 in the Sindhudurg district of Maharashtra state in Western India, which coincides with cashew flowering in the region. It is situated in the northern part of the Western Ghats-Sri Lanka Biodiversity Hotspot. The average annual rainfall is 3,500 mm, and the annual temperatures range between 12°C to 40°C (Munje & Kumar, 2022). Most sites were located at low elevations (not exceeding 600 m.a.s.l). The landscape primarily consists of moist deciduous and evergreen forest patches, which host ornithophilous plant species such as *Canthium rheedei*, *Ixora brachiata*, and *Lophopetalum wightianum*, among others. Please see Madhu et al., (2025) for additional information on study area.

Most of the forests in the low-elevation areas of northern Western Ghats, which are suitable for cashew cultivation, are privately owned. Protected Areas cover less than 1.5% of the geographic area of the low-elevation forests of northern Western Ghats of Maharashtra state. Over the last two decades, the area has witnessed a rapid conversion of these privately-owned forests to cashew plantations (Rege & Lee, 2022). More than a third of the two district sub-divisions are under cashew plantations. While cashew was historically grown as agroforests with cashew trees planted amidst native trees, today cashew is grown as monocultures. In this region, Cashew flowers between the end of December to March with peak flowering in the end of January and early February. Key nectar resources for birds in these modified systems include *Helicteres isora*, *Moullava spicata*, and *Grewia tiliifolia*, among others (Mahale, 2023).

### 3.2. Field Methods

#### 3.2.1. Bird sampling

To estimate the diversity and abundance of nectar-feeding birds across land-use types, NM and VS conducted variable-radius point-count surveys at 100 locations. Points were distributed across cashew monocultures (n=50), cashew agroforests (n=25), and forests (n=25) (Fig. S1), with each point separated by at least 250 m to minimise spatial autocorrelation (Lee et al., 2024; Madhu et al., 2025). We visited each point five times between January and April, 2024. Sampling was carried out in the mornings between 0645 and 1000 hr. At each visit, 10-minute point-counts were conducted, during which species identity, detection distance, and the number of individuals seen or heard (excluding flyovers) were recorded. Bird abundance data were pooled across the five replicates for all bird-related analyses (Shahabuddin et al., 2021). Only individuals detected within 100 m of the observation point were included in the analysis. Distances of birds from the observation point were recorded as whole numbers (when detected visually) and in the following radial distance bins (in m) (when detected auditorily): 0–5, 6–10, 11–15, 16–20, 21–30, 31–50, 51–75, and 76–100.

#### 3.2.2. Floral availability

At each bird observation point, we established a 10-m radius circular plot to quantify floral resources. Within each plot, we counted the total number of open floral units and the species identity of flowering plants. To account for seasonal variation in resource availability, we sampled each point five times between January and April, coinciding with bird surveys. For the estimation of floral abundance and diversity, we included only those plant species known to provide resources to nectarivorous birds (Mahale, 2023; Sadekar, 2024) and for which we observed bird-flower interactions.

#### 3.2.3. Visitation data

We recorded bird visits to flowering plants in forests, cashew agroforests, and cashew monocultures between January and May 2024. Observations were conducted both during point counts and time-constrained searches. The searches were conducted, between 0600 to 1800 hr and the total sampling effort for searches was 52 hours (Table S1). We recorded every individual observation of a bird probing or predating on the flower as an interaction. We classified each interaction as either pollination, nectar-robbing (when the bird pierced the flower’s corolla at the base to access the nectar), or predation (when the birds plucked the flower and swallowed it). Predation events were excluded from all analyses. We included visits observed on the edges of monoculture plantations in all analyses.

### 3.3. Analysis

We conducted all the analyses in R version 4.4.0 (R Core Team, 2024).

#### 3.3.1. Abundance of flowers

To determine the effects of land-use type and time (since flower availability changed from January to April) on flower abundance, we used a generalised linear mixed model (GLMM) with a zero-inflated negative-binomial error structure. A zero-inflated model was appropriate as forest data contained a high proportion of zeros (Table S2). We used a random intercept model with point ID as a random effect and the interaction between day (Julian day of the year) and land-use type as the fixed effect. We considered predictor variables to significantly influence flower abundance if their beta coefficients had 95% confidence intervals (CIs) that did not overlap zero. We also estimated marginal and conditional *R^2^* values for the model. We used R packages ‘glmmTMB’, ‘DHARMa’, ‘performance’, and ‘MUMIn’, for this analysis (Brooks et al., 2017; Hartig, 2016; Kamil Bartoń, 2010; Lüdecke et al., 2021).

#### 3.3.2. Abundance of nectar-feeding birds

We estimated the density of nectar-feeding birds using the multi-covariate distance sampling (MCDS) approach in the R-package ‘Distance’ (version 2.0.0, Miller et al., 2019). We estimated densities for nine species of nectar-feeding birds that were detected at least ten times during point counts and recorded in at least ten bird-flower interactions (Table S3 and S4). The remaining species were extremely rare and were therefore excluded, as they were unlikely to influence the bird densities and interpretation of results. We truncated detections beyond 50 m from the point of observation to improve model fit. We fitted half-normal and hazard-rate detection function models and selected the best-fitting model based on the lowest Akaike Information Criterion value (Buckland et al., 2015). Densities were estimated separately for each land-use type.

#### 3.3.3. Diversity of flowers, nectar-feeding birds, and bird-flower interactions

To determine differences in the diversity of flowers, nectar-feeding birds, and their interactions across land-use types, we used the Hill numbers framework of diversity measures, implemented through a sample-coverage-based rarefaction approach (Roswell et al., 2021). We estimated species richness (q = 0), and Hill-Shannon diversity (q = 1) based on abundance data. Species richness is highly sensitive to rare species, while Hill-Shannon incorporates both rare and common species, weighting them by their relative abundances in the community. For flowers, we pooled data across five replicates for 18 tree species identified as nectar resources for birds based on literature and observed interactions. For birds, we pooled data across five replicates for 28 species recorded during point counts and observed visiting open flowers. For interactions, we considered only those visits where an individual was observed to be feeding on nectar and excluded flower predation events. Interaction diversity, estimated from interaction frequencies, is influenced by both the number of interacting species and the distribution of interaction weights among them (García, 2016). We assigned each unique interaction a separate identity (Dyer et al., 2010), equivalent to a “species” in flower and nectar-feeding bird diversity. We used the R package ‘iNEXT’ to estimate sample coverage and Hill-Shannon diversity (Chiu et al., 2023; Hsieh et al., 2015).

#### 3.3.4. Composition of flowers and nectar-feeding birds

We examined differences in composition of flowers and nectar-feeding birds across land-use types (forest, agroforest, monoculture) using non-metric multidimensional scaling (NMDS) with the Bray-Curtis dissimilarity index. We log-transformed the flower abundance data due to the inflated presence of zeros. We performed NMDS on abundance data pooled across temporal replicates for both flowers and nectar-feeding birds. Since the stress values of the NMDS for nectar-feeding birds was greater than 0.2, we used ecological null models using the ‘swap-count’ method following Dexter et al., (2018) to validate the NMDS ordination fit. We performed an analysis of similarities (ANOSIM) to test for significant differences in flower and nectar-feeding bird composition between land-use types. Additionally, we conducted the Mantel’s test with 1,000 permutations to determine correlation between the dissimilarity matrices of flower and nectar-feeding bird compositions. We used functions *metaMDS*, *oecosimu*, *anosim*, and *mantel* from R-package ‘vegan’ to conduct these analyses (Oksanen et al., 2001).

#### 3.3.5. Network organisation

We compared the organisation of plant and nectar-feeding bird communities across land-use types by constructing weighted, quantitative bipartite networks for each land-use category. The numbers of bird visitations to flower species were used as weights. We estimated widely established network-level metrics: connectance, complementary specialisation (*H_2_’*), weighted Nestedness Overlap and Decreasing Fill (*wNODF*), and modularity (*Q*) for each land-use type separately. We compared the observed values of these metrics to the means of 1,000 null model networks generated using the Patefield algorithm that preserves marginal totals of the observed interaction matrix. Rarefaction has been recommended in cases where network sizes are variable (Blüthgen & Staab, 2024), since certain network metrics (e.g., connectance, modularity) are sensitive to network size (Vanbergen et al., 2017). In our case, network sizes differed across different land uses (forest: 171, agroforest: 216 and monoculture: 243). Therefore, we rarefied each network 100 times by standardising total interaction frequency to the lowest number of interactions (171 interactions observed in forests) observed across land-use types. On these rarefied networks, we recalculated connectance, complementary specialisation, wNODF, and modularity. We then obtained the mean and standard deviation (SD) of each metric from the 100 rarefied subsamples per land-use type. This allowed us to determine whether the observed metric for forests fell within 2 SD units of the estimated metrics for agroforests and monoculture plantations. All analyses were performed using ‘bipartite’, ‘tidyverse’, and ‘purr’ packages in R (Dormann et al., 2008).

## 4. RESULTS

### 4.1. Abundance of flowers

We estimated flower abundance of 18 flowering plant species belonging to 11 plant families (Table S2). Open flowers were dominated by non-native species such as cashew (*Anacardium occidentale*) in agroforests and monocultures, and native species such as *Terminalia paniculata* in forests and agroforests, and *Grewia tiliifolia* in agroforests (Fig. S2). Peak flower abundance was observed in January and February (Fig. 1a). We found statistically supported interaction effects between land-use and Julian day, indicating that the rate of decline in flower abundance over time differed across land-use types (Fig. 1a, Table S5, *R^2^_marginal_* = 0.7). While the flower abundance was higher in monoculture and agroforests as compared to forests, so was the rate of decline in flower abundance over time (*p* < 0.001) (Fig. 1a, Table S5).

**Figure 1.**
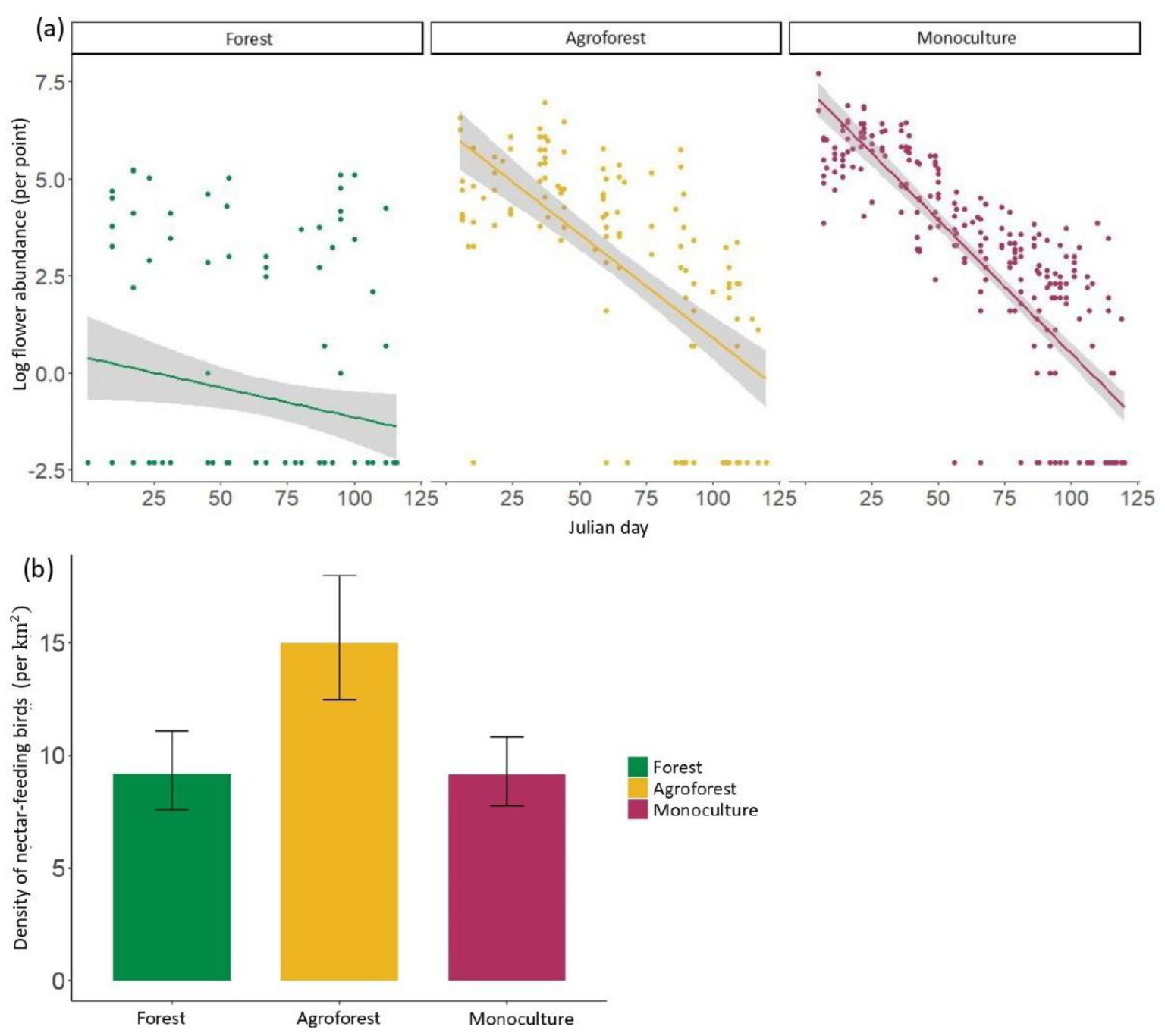
Plots showing (a) the interactive effect of the Julian day and land-use type on log-transformed flower abundance per plot across the three land-use types. Solid lines represent best-fit lines from a standard regression model and shaded regions represent associated 95% CIs. (b) Estimated densities of nectar-feeding birds across land-use types. Error bars represent 95% CIs.

### 4.2. Density of nectar-feeding birds

In our 10-minute variable-radius point counts, we recorded 4,111 detections of 28 out of the 31 bird species that we observed interacting with flowers (Table S3). Density (± SE) of nectar-feeding birds was highest in agroforests (15.0 ± 1.4 flocks per km^2^) (Fig. 1b). Forests and monocultures had similar densities (Forest: 9.2 ± 0.9 flocks per km^2^, Monoculture: 9.2 ± 0.8 flocks per km^2^) (Table S7 and S8).

### 4.3. Diversity of flowers, nectar-feeding birds, and nectarivorous interactions

In total, we documented 632 interactions between 31 bird species and 45 flower species. Of these, we could confirm 15 nectar-robbing interactions, and one instance each of *Ocyceros griseus* and *Oriolus kundoo* consuming the flowers of *Carallia brachiata*. For the remaining 615 interactions, birds probed the flower without visible damage or nectar theft. Among birds, 274 interactions were of the endemic *Leptocoma minima,* followed by 122 interactions of wide-ranging *Cinnyris asiaticus* (Table S4). Among plants, 204 interactions were for *Anacardium occidentale* (cashew), followed by 94 interactions on *Getonia floribunda* (Table S6). We detected 171 interactions in forests, 216 in agroforests, and 243 in monocultures.

The minimum sample coverage for the flower and bird abundance data was very high (0.99). However, for the interaction data, the sample coverage for forests was 0.71, while that for agroforest and monoculture was 0.80 and 0.79, respectively. Flower species richness (q = 0) was 6.5 times higher in forests and 5.2 times higher in agroforests than in monocultures (Table S9, Fig. 2a). Similarly, Hill-Shannon diversity (q = 1) of flowers was 3.5 times higher in forests and ∼ 3.2 times higher in agroforests than in monocultures (Table S10, Fig. 2a). Agroforests had the highest species richness (q = 0) and Hill-Shannon (q = 1) diversity of nectar-feeding birds across land-use types (Table S11 and S12, Fig. 2b). In contrast, forests had the highest mean richness (q = 0) and Hill-Shannon (q = 1) diversity of interactions across land-use types (Table S13 and S14, Fig. 2c).

**Figure 2.**
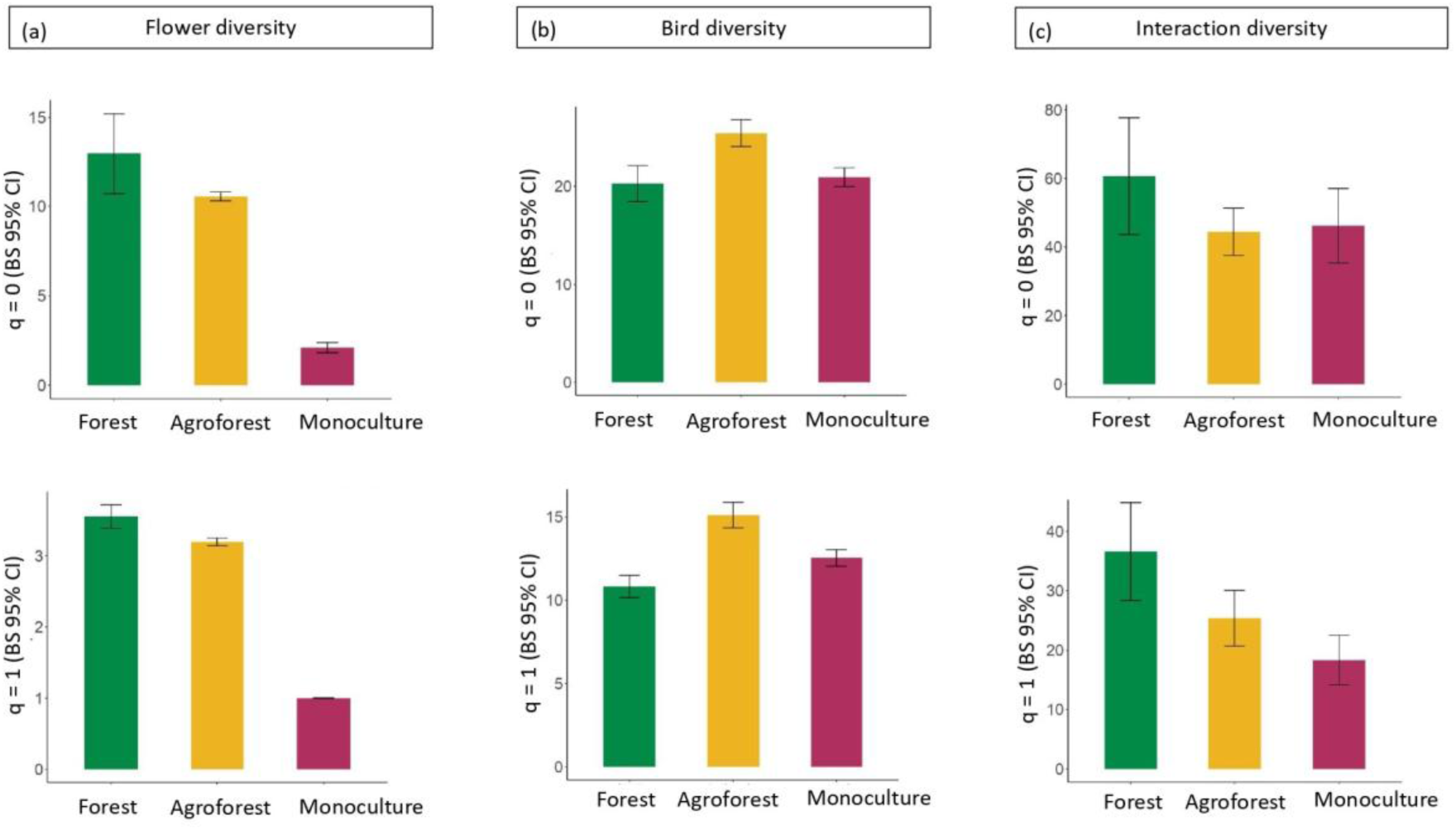
Hill species richness (q = 0), and Hill-Shannon (q = 1) of (a) flower diversity per plot, (b) nectar-feeding bird diversity per plot, and (c) bird-flower interaction diversity across the three land-use types. Error bars represent bootstrapped (n = 1000) 95% CIs.

### 4.4. Composition of flowers and nectar-feeding birds

Floral communities differed significantly across land-use types (*R_anosim_* = 0.69, *p* = 0.001, Stress = 0.11; Fig. 3a). Similarly, the composition of nectar-feeding bird species also differed significantly across land-use types (*R_anosim_* = 0.5, *p* = 0.001, Stress = 0.2; Fig. 3b). The null hypothesis for no difference in bird composition was rejected by a one-sample *z*-test after 1,000 permutations, indicating significant differences in community composition (*z* = −2.62, *p* = 0.003). Moreover, the flower and bird compositional distance matrices were significantly correlated (Mantel’s *r* statistic = 0.53, *p* < 0.01), indicating that sites differing in flower composition also exhibited corresponding differences in nectar-feeding bird communities.

**Figure 3.**
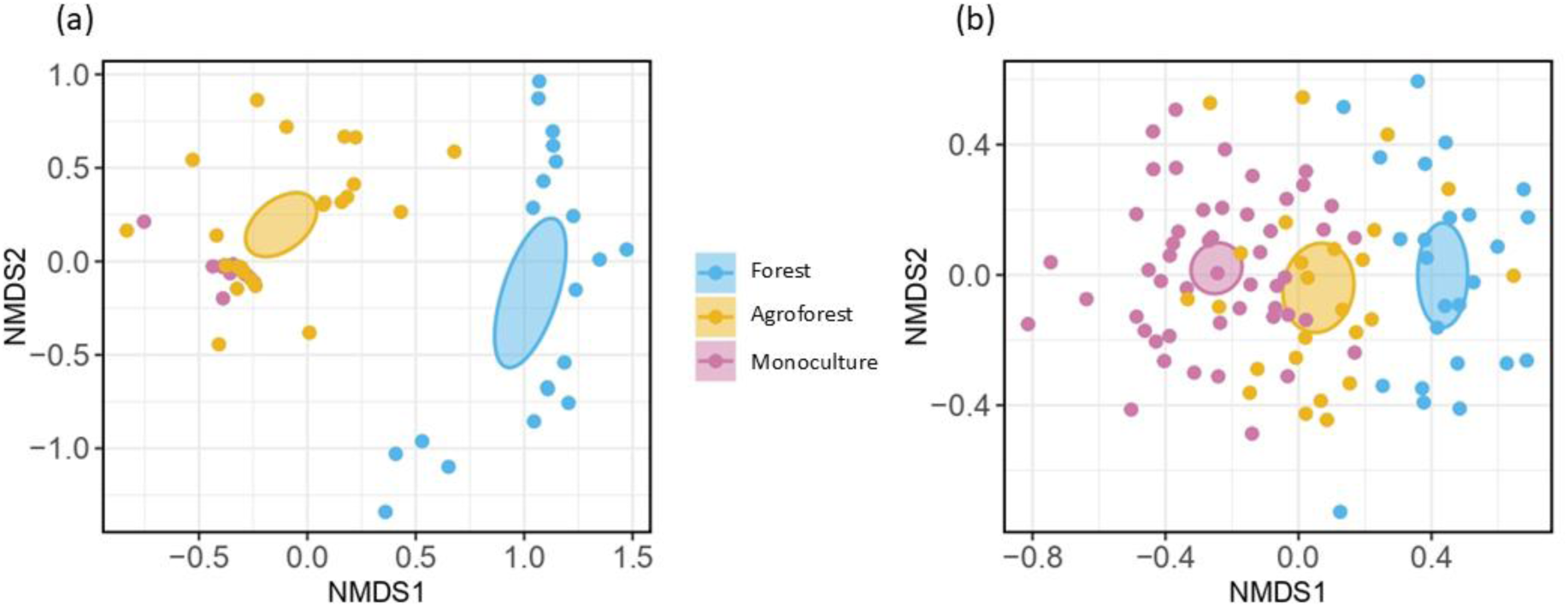
Non-metric multidimensional scaling ordinations showing differences in (a) ornithophilous flower species composition, and (b) nectar-feeding bird species composition across the three land-use types. Ellipses represent 95% confidence intervals around group centroids.

### 4.5. Network organisation

Networks in all three land-use types (forests, agroforests, and monoculture plantations) were significantly less connected and more specialised and modular than their respective null models (Table 1). However, only networks in agroforests and monocultures were significantly less nested than expected under the null models (Table 1). After rarefying networks in agroforests and monocultures to match the size of the forest network (100 iterations), the observed forest network showed lower connectance, specialisation, and modularity (Table S15, Fig. S4). These values fell outside the 95% confidence intervals (± 2 SD) of the rarefied agroforest and monoculture networks (Fig. 4), suggesting consistent structural differences in network organisation across land-use types, even when controlling for network size.

**Figure 4.**
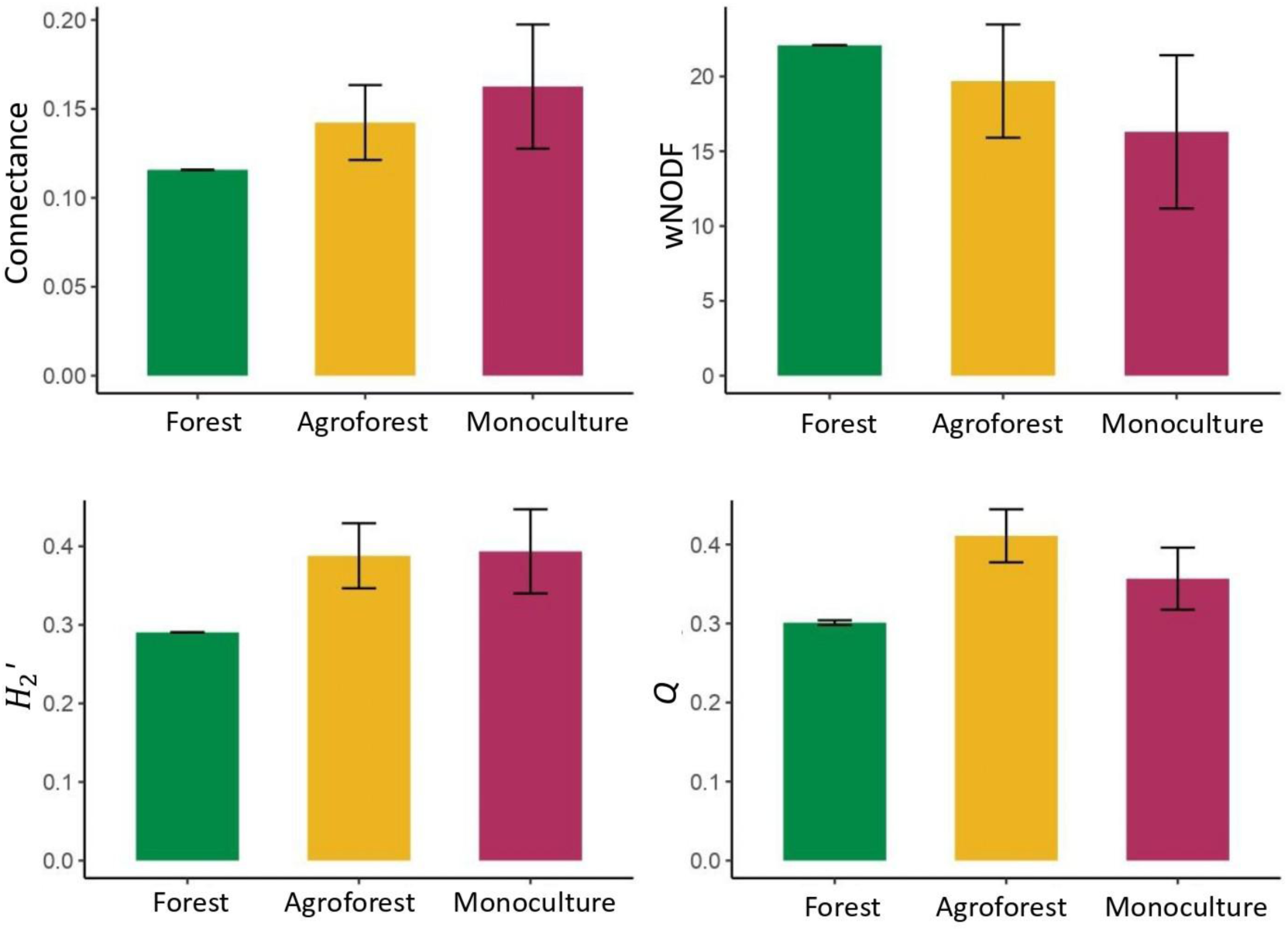
Network parameters (Mean ± 2 SD) across land-use types after rarefying agroforest (size: 216) and monoculture plantation (size: 243) networks to the size of forest network (size: 171). We rarefied by subsampling the agroforest and monoculture network 100 times.

**Table 1.** Observed network parameters for forests, agroforests, and monocultures in the northern Western Ghats, India. Asterisks indicate values that were significantly different (*p* < 0.05) from 1,000 null models. Visualisations of these networks have been provided in Fig. S3.

| Parameter | Forest | Agroforest | Monoculture |
| --- | --- | --- | --- |
| Connectance | 0.116* | 0.151* | 0.159* |
| z-score | -5.616 | -7.974 | -2.921 |
| $H_2'$ | 0.290* | 0.377* | 0.361* |
| z-score | 3.916 | 13.675 | 11.704 |
| wNODF | 22.078 | 22.826* | 17.971* |
| z-score | -1.664 | -3.214 | -5.341 |
| Modularity (Q) | 0.297* | 0.402* | 0.151* |
| z-score | 4.232 | 13.607 | 13.26 |

## 5. DISCUSSION

We found that forests, cashew agroforests, and monocultures hosted distinct communities of flowers and nectar-feeding birds. Resource abundance decreased most rapidly in monocultures over time, followed by agroforests. Agroforests had a significantly higher density of nectar-feeding birds than forests and monocultures. Diversity of flowers and interactions were significantly higher in forests as compared to agroforests and monocultures. However, the diversity of nectar-feeding birds was highest in agroforests. Networks in all land-use types were significantly less connected and more specialised and modular than expected by chance. However, only agroforests and monocultures had significantly more nested networks compared to null models. After rarefaction, the forest network was less connected, specialised and modular than those in agroforests and monocultures. Our findings highlight the negative impacts of agricultural intensification on the diversity and abundance of nectar resources and nectar-feeding birds, and consequently in bird-flower interactions and network organisation.

### 5.1. Flower abundance and diversity across land-use types

Modified habitats such as agroforests and monoculture plantations offered seasonally abundant but less diverse and compositionally distinct flower resources compared to forest habitats. Existing literature on flower diversity and abundance in modified landscapes vis-à-vis natural habitats mostly comes from grassland ecosystems in temperate regions (Neumüller et al., 2020; Potts et al., 2004; Weiner et al., 2011). Published information on flower availability in tree crops and tropical systems is scarce (but see Eeraerts et al., 2021; Klein et al., 2002). In line with previous studies that report higher diversity of flower resources in more natural habitats (Holzschuh et al., 2008; López-Flores et al., 2024), we found that flower richness was the highest in forests, followed by agroforests.

Unlike flower richness that is often found to be higher in natural conditions (López-Vázquez et al., 2024; Papanikolaou et al., 2017; Wray et al., 2014), flower abundance can be higher (Carman & Jenkins, 2016), lower (Adedoja & Kehinde, 2018), or show no difference (Durán & Kattan, 2005) in modified habitats compared to natural areas. Cashew agroforests and monoculture plantations exhibit temporal discontinuity, with a period of hyperabundance during flowering, followed by little to no provision for the remaining months (Saunders et al., 2015). As time progressed, the decline in flower abundance was the steepest in monocultures, mirroring findings from farmland ecosystems (Timberlake et al., 2019, 2021). While a previous study highlighted the importance of native trees in plantations for evolutionarily distinct bird species (Madhu et al., 2025), our findings demonstrate their additional role as key nectar sources for flower-visiting birds (Fig. S5). Planting zoophilous native plants that bloom during the cashew off-season along the boundaries of monocultures can potentially enhance their suitability for nectar-feeding fauna. Native plants along field edges were found to attract invertebrate flower-visitors such as bees in coffee plantations in Brazil (Pereira Machado et al., 2024) and cucumber fields in Wisconsin (Lowe et al., 2024). While seasonally, monocultures may offer abundant flower resources, given the lower diversity of flowers in modified ecosystems, it may compromise the nutritional and sugar composition of nectar available to nectar-feeding fauna (Inês Da Silva et al., 2024; Jones & Rader, 2022; Pioltelli et al., 2024). These alterations would likely have negative implications for bird physiology (Nicolson & Fleming, 2003), an aspect that is relatively understudied.

### 5.2. Abundance and diversity of nectar-feeding birds across land-use types

Nectar-feeding birds are known to benefit from certain levels of deforestation, particularly in agroforestry context (Tscharntke et al., 2008). Species richness and abundance tend to either increase (Davies et al., 2015; Waltert et al., 2004) or remain the same (Bustamante-Castillo et al., 2020; Durán & Kattan, 2005) with agricultural intensification. Intermediate land uses such as agroforestry can balance the tradeoffs between agriculture and biodiversity and have been proven successful in sustaining nectar-feeding bird populations (López-Flores et al., 2024). In line with these results, the relatively less-disturbed cashew agroforests in our study had the highest diversity and abundance of flower-visiting birds. Monocultures had absolute richness (q = 0) comparable to forests, and Hill-Shannon diversity (q=1), which emphasizes common nectar-feeding bird species, was higher than in forests. However, the composition of nectar-feeding bird species was different between forests and cashew plantations likely caused by filtering out of forest specialist nectar-feeding birds in cashew plantations as reported elsewhere (Infante et al., 2020). Past research in the landscape (Madhu et al., 2025) has demonstrated variable responses of nectar-feeding birds to forest conversion. For example, the Nilgiri Flowerpecker *Dicaeum concolor* and Little Spiderhunter *Arachnothera longirostra* responded positively to native trees within cashew plantations, whereas Ashy Prinia *Prinia socialis* and Grey-breasted Prinia *Prinia hodgsonii* responding negatively. Agroforests hosted a nectar-feeding bird composition in between that of forests and monocultures, implying the presence of both forest specialist as well as disturbance-adapted species.

The endemic Crimson-backed Sunbird *Leptocoma minima* and the widespread Purple Sunbird *Cinnyris asiaticus* participated in the highest number of interactions and visited the highest number of plant species. They also exhibited high visitation frequencies on individual flowers (NM, VS, pers. obs.), showcasing their important role as pollinators in the region.

### 5.3. Interactions and network indices across land-use types

Agroforests, despite having higher diversity of nectar-feeding birds as compared to forests, had lower interaction diversity. This suggests that interaction diversity may have been strongly influenced by flower diversity rather than by nectar-feeding bird diversity. Monoculture plantations often had flowering species planted along their edges such as *Helicteres isora* and the non-native *Gliricidia sepium* that attracted numerous visitors while in bloom. Hence, although floral richness was minimal, these plantations still hosted interactions among various plant-bird species pairs, contributing to overall interaction diversity.

In contrast to results from other studies, the forest community in our study was less specialised and modular compared to those from agroforest and monoculture plantations (Lázaro et al., 2016; Morrison & Mendenhall, 2020). However, connectance was lowest in forests, mirroring the findings of Morrison et al., (2020) and Shinohara et al., (2019) who found higher connectance in disturbed or modified habitats. These patterns are likely an outcome of the high abundance of cashew flowers as compared to other species in agroforests and monocultures (Fig. S2). Most visits in plantations were to cashew flowers, despite the presence of other species in the interior or along plantation edges. The high availability or nectar quality of cashew could have attracted numerous nectar-feeding birds, maintaining specialised and compartmentalised communities centered around it (Watts et al., 2016). In parallel, these nectar-feeding birds often included generalist feeders such as Purple Sunbird *Cinnyris asiaticus* that fed on flowers of other plant species as well, thus maintaining high connectance within the networks.

### 5.4. Caveats and way forward

One may argue that our networks were constrained by limited sampling completeness, particularly in forests. To address this and allow for suitable comparisons of network parameters across land-use types, we rarefied the agroforest and monoculture interaction networks to the network size of forests. Since the study was focussed on comparing flower and nectar-feeding bird interactions, we conducted the study during the flowering period of cashew in the region, which also happens to coincide with the peak flowering of several shrub and tree species in the Western Ghats (Bhat & Murali, 2001; Sundarapandian et al., 2005). Future work could assess the variation in nectar quality and volume of flowers in monocultures, agroforests, and forests and determine their influence on nectar-feeding birds. It would be interesting to assess how sunbirds utilise resources in cashew agroforests during the non-flowering period of cashew, as they may switch to invertebrate diets during this time (Sadekar, 2024). Twelve of the 32 nectar-feeding bird species documented in this study were observed feeding on cashew flowers. Given that the flowers are small, and the birds were not observed robbing the nectar of cashew (Satish et al., in prep.), most species likely function as pollinators for cashew. Future studies should assess whether proximity to forests influences the abundance of nectar-feeding birds in cashew agroforests and whether this translates into greater pollination services by measuring fruit set after accounting for contributions made by insect pollinators. Such studies could potentially help inform management recommendations, including planting native trees in cashew plantations to enhance biodiversity and benefit local cashew farmers (Kaiser-Bunbury & Blüthgen, 2015; Peters et al., 2016).

## 7. CRediT AUTHORSHIP CONTRIBUTION STATEMENT

Conceptualisation: RN, NM, RJ; Data curation: NM, VS; Formal analysis: NM, RN, RJ; Funding acquisition: NM, RN; Investigation: NM, VS; Methodology: RN, NM, RJ; Project administration: VS, NM; Resources: NM, RN; Supervision: RN, RJ; Visualisation: NM, RN; Writing – original draft: NM, RN; Writing – review & editing: RJ, VS. All authors in this study are indigenous to the country where the study was conducted. Whenever relevant, literature published by scientists from the region has been cited.

## 8. ACKNOWLEDGEMENTS

We thank the Dean and Director of the Wildlife Institute of India, and the Maharashtra Forest Department for giving us the necessary permission (No: Desk-22(8)/WL/Research/C. No. 103/635/24–25) to conduct the study. We thank the Wildlife Institute of India, Godrej Consumer Products Limited and Rohini Nilekani Philanthropies for funding this study. We thank Vedika Dutta and Rintu Mandal for help with analyses, and Navendu Page for inputs and help with plant identification. We thank Anand Osuri for his valuable feedback. We are grateful to Praveen and Shital Desai and Vanoshi Forest Homestay for support during fieldwork. We thank cashew landowners in Dodamarg who allowed us to conduct fieldwork on their properties.

## 10. DATA AVAILABILITY

The data and code that support the findings of this study will be made openly available on Zenodo upon acceptance of the manuscript.

## SUPPLEMENTARY INFORMATION

### Supplementary figures

**Figure S1.**
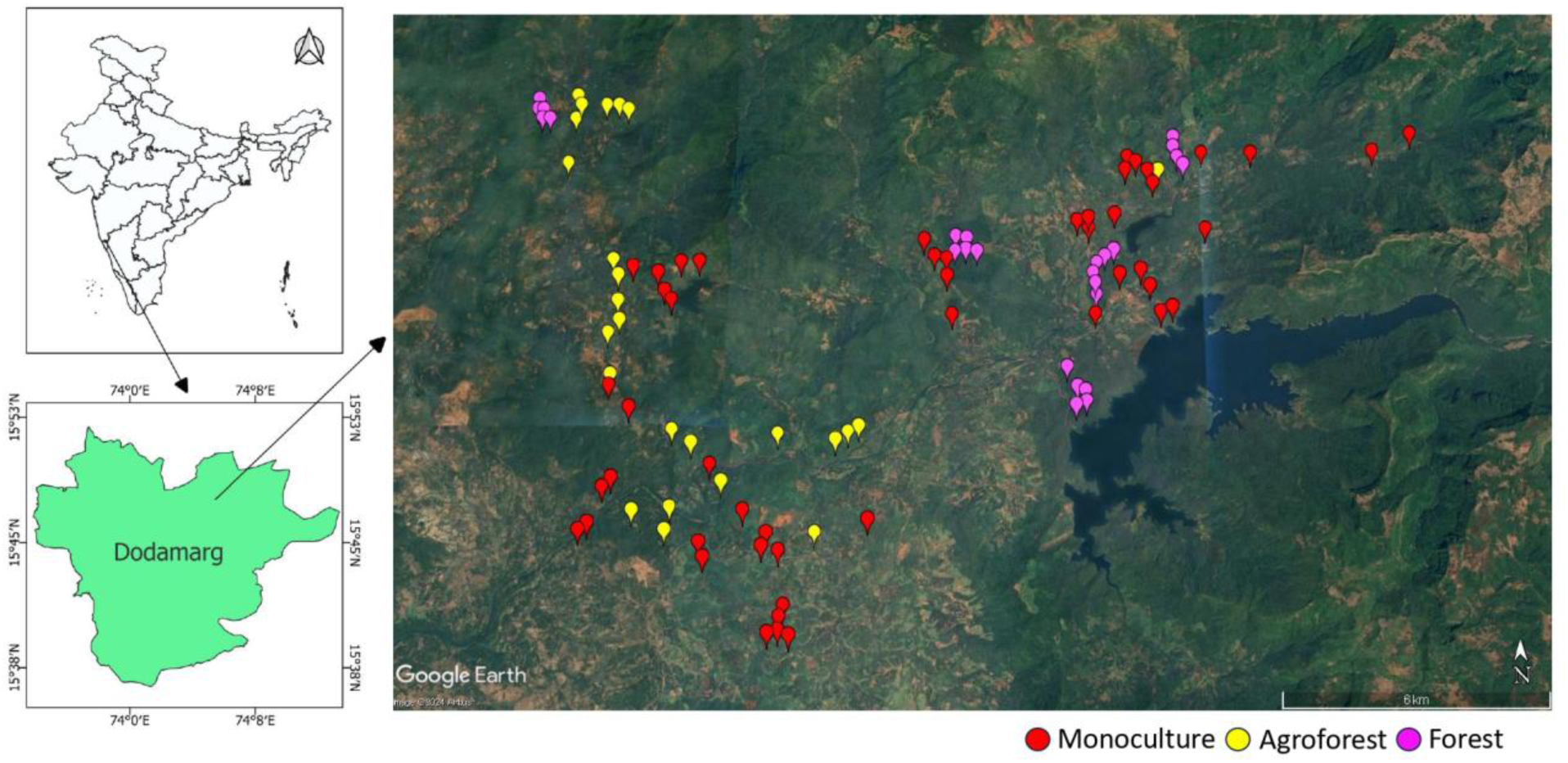
The satellite image displays 100 sampling points (monocultures, agroforests, and forests) in Dodamarg Taluka of Sindhudurg district, India. These points were used to estimate the diversity of nectarivorous birds across land-use types.

**Figure S2.**
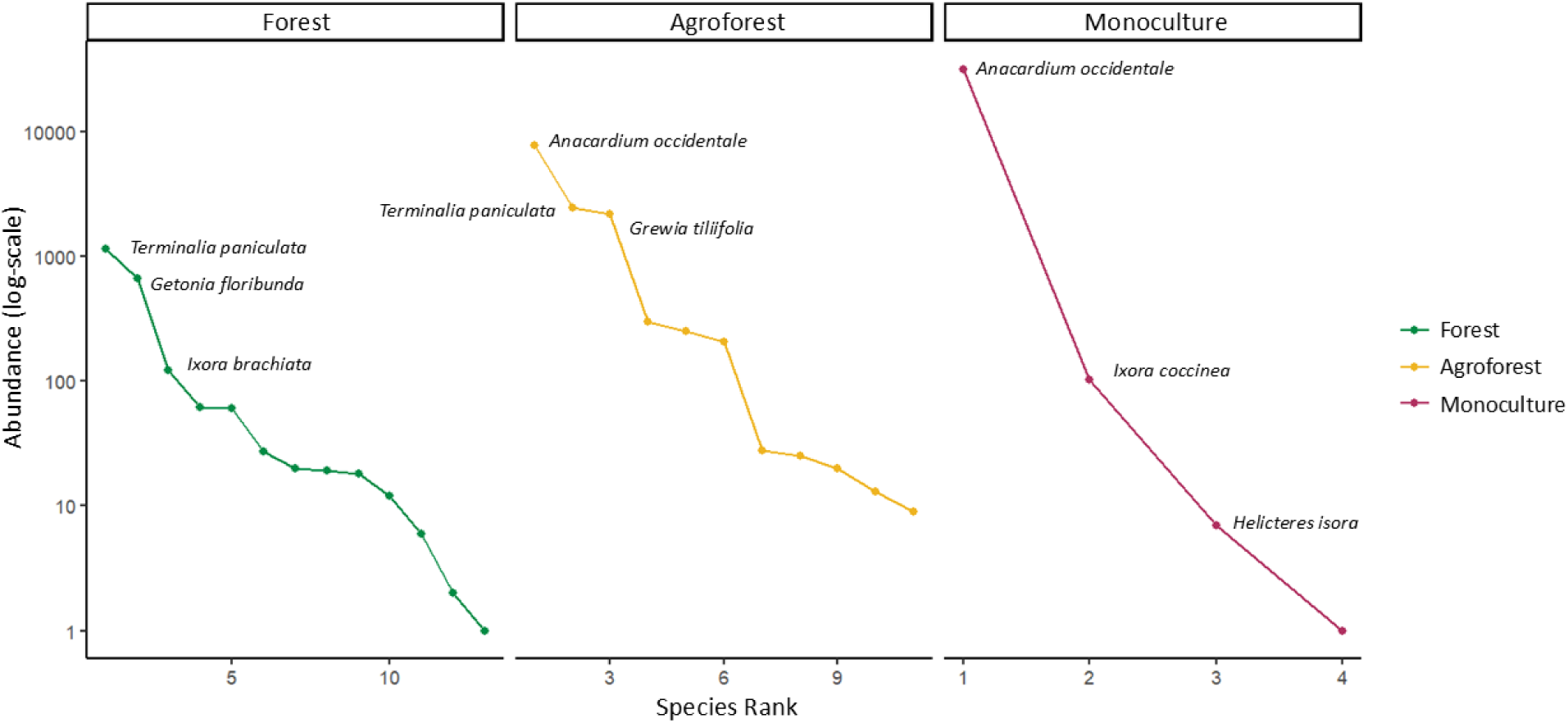
Rank-abundance curves of flowering species across land-use types. The top three ranked species are labeled.

**Figure S3.**
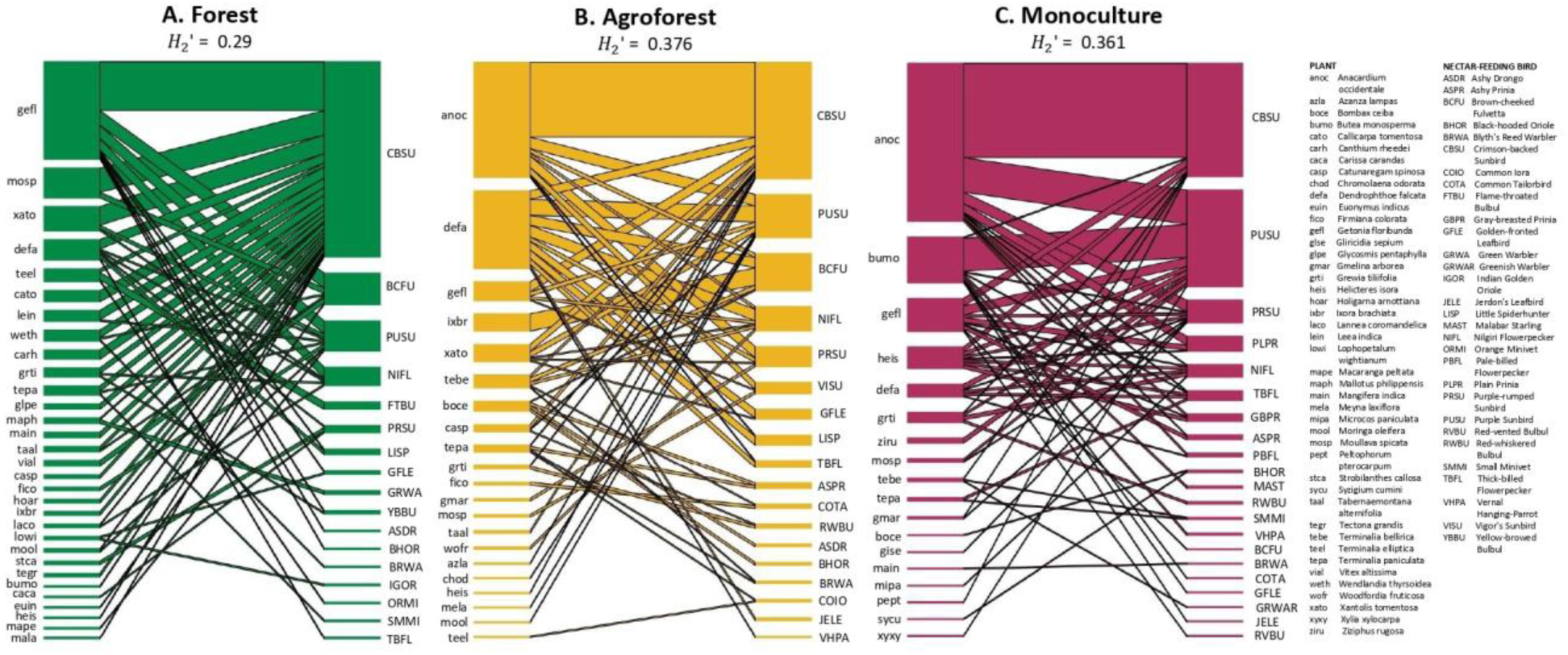
Plant-nectar-feeding bird networks across land-use types and their specialisation (H_2_’) values. Boxes on the left of each network represent plant species and those on the right represent bird species. Heights of boxes represent relative participation frequencies of plant and bird species in each network. The width of a connecting line between a pair of plant and bird species represents the interaction frequency between them. Plant and bird species are placed in the decreasing order of the row and column totals in the input interaction matrix. Code names and common names of plant and bird species are mentioned alphabetically. Scientific names of bird species (Jetz et al., 2012) are in Table S4.

**Figure S4.**
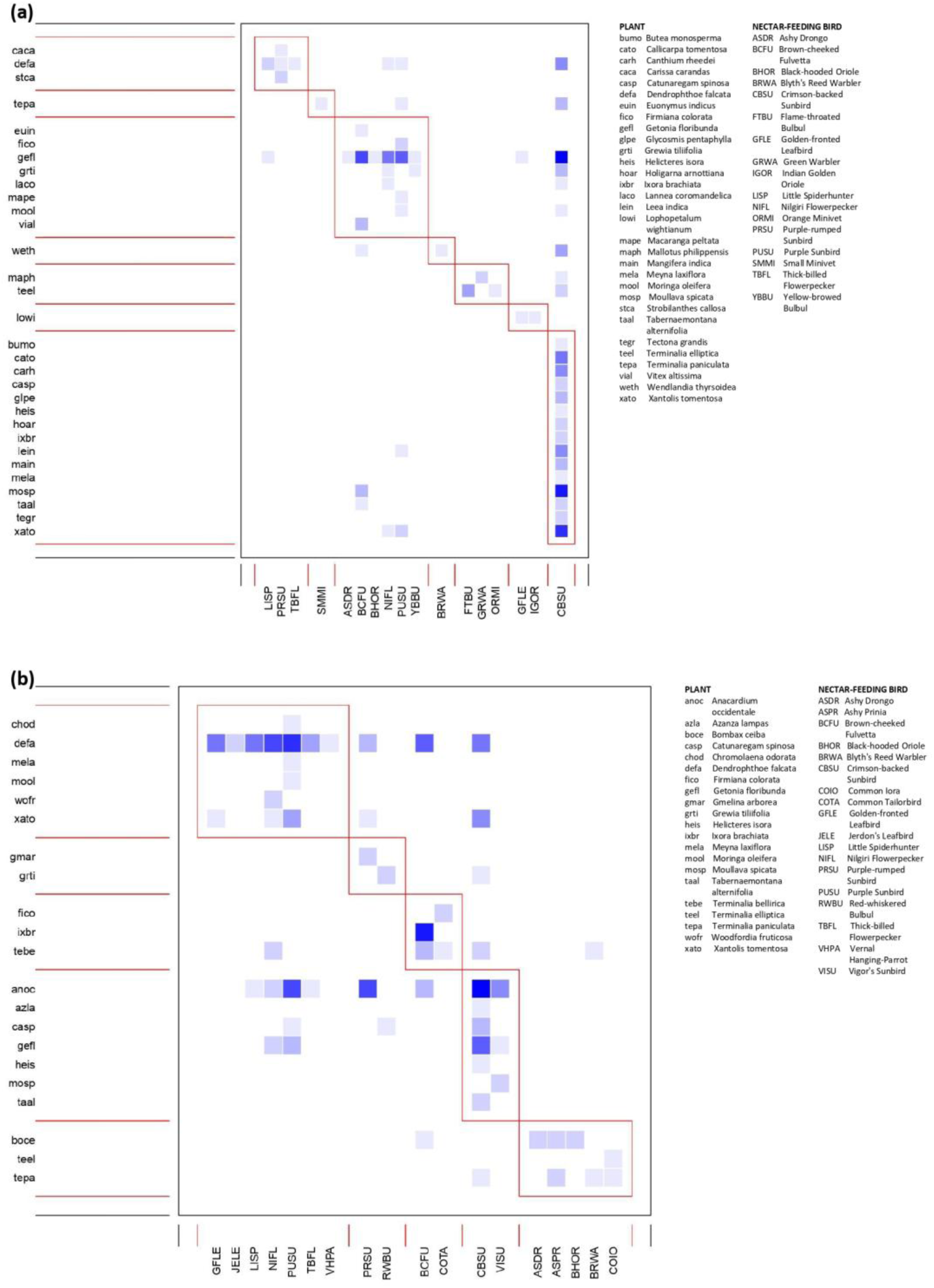
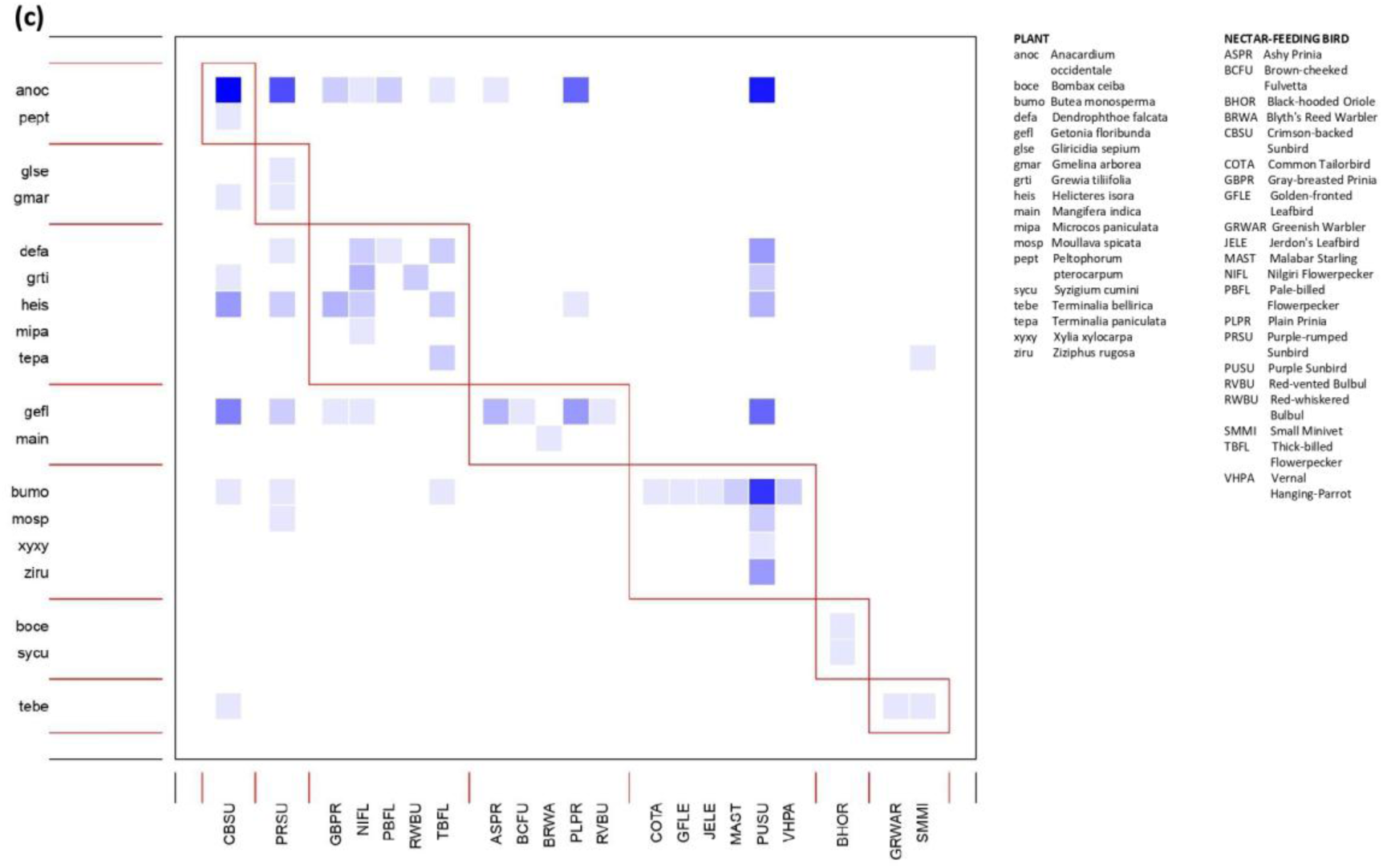
Modules of interaction networks computed for (a) forest, (b) agroforest, and (c) monoculture, using the *computeModules* function of the ‘bipartite’ package in R.

**Figure S5.**
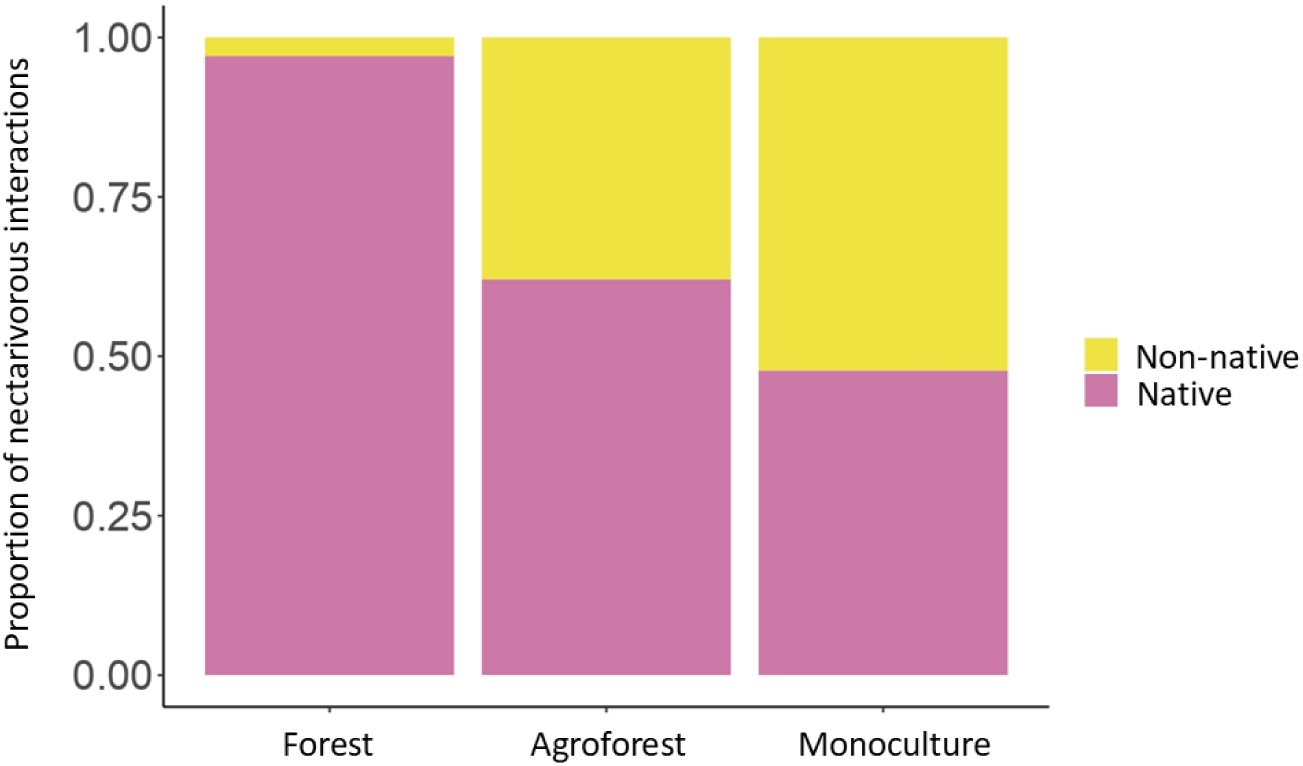
Stacked plot displaying the proportion of interactions between nectar-feeding birds and native and non-native plants across the three land-use types (forest, agroforest, and monoculture).

### Supplementary Tables

**Table S1.**
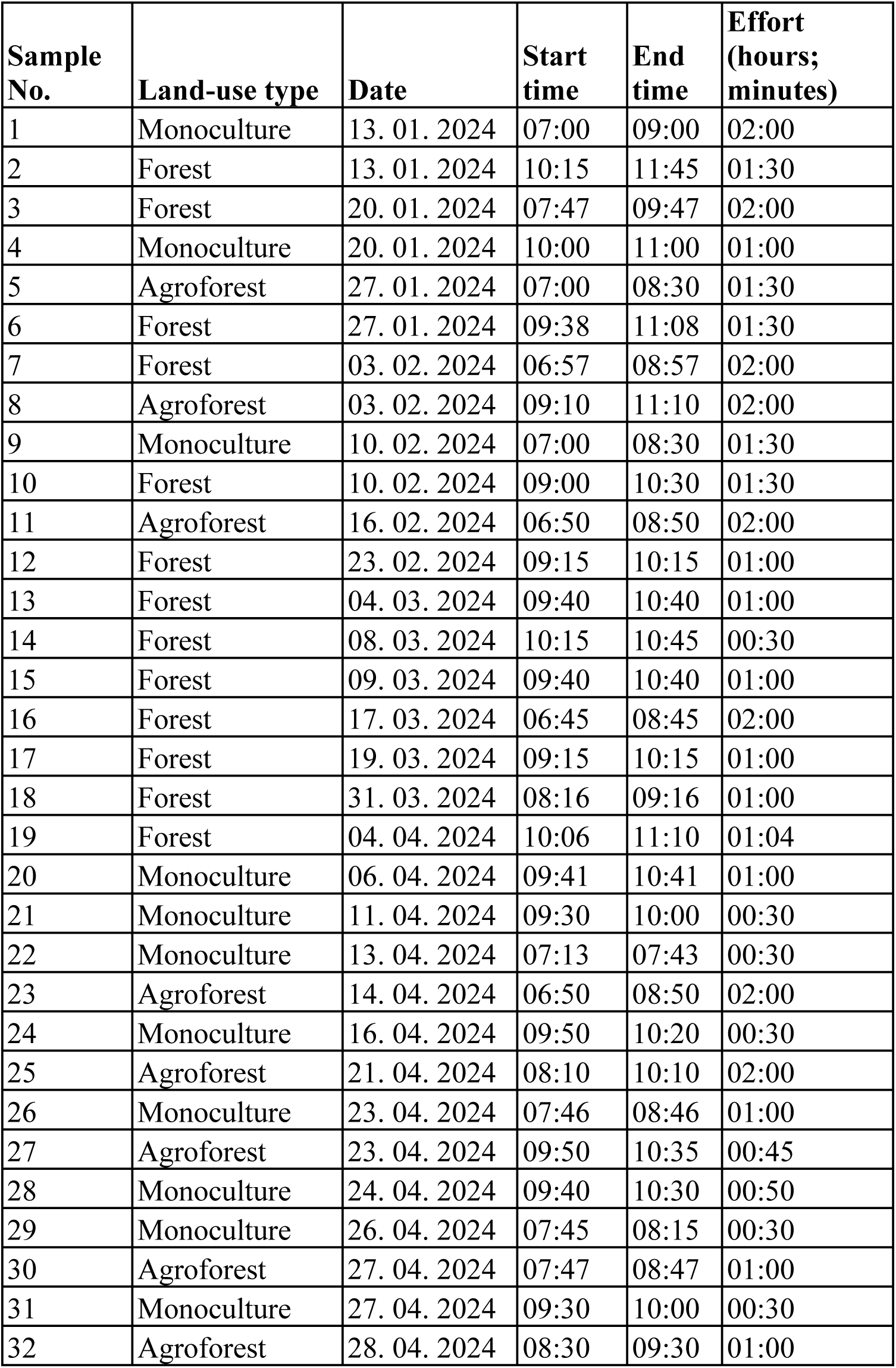
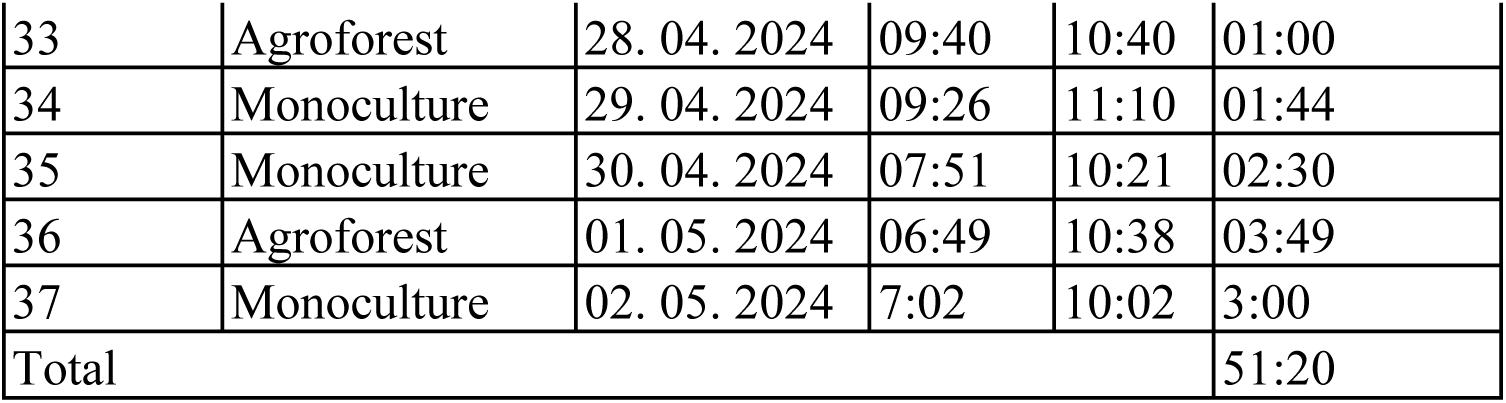
Sampling effort of interactions across forests, agroforests, and monocultures along with dates, start-times, and end-times.

**Table S2.** Checklist of plant species that flowered during the sampling period, either known to be resources to birds or visited by birds during the sampling period, and the abundance of flowers they hosted within 10 m circular radius plots across 25 forest points, 25 cashew agroforest points, and 50 cashew monoculture points. Species are sorted according to families.

| Sl. No. | Family | Species | Total abundance of open flowering units across forest plots | Total abundance of open flowering units across agroforest plots | Total abundance of open flowering units across monoculture plots |
| --- | --- | --- | --- | --- | --- |
| 1 | Anacardiaceae | *Anacardium occidentale* | 0 | 7787 | 31808 |
| 2 | Apocynaceae | *Tabernaemontana alternifolia* | 62 | 13 | 1 |
| 3 | Combretaceae | *Getonia floribunda* | 661 | 207 | 0 |
| 4 | Combretaceae | *Terminalia elliptica* | 20 | 20 | 0 |
| 5 | Combretaceae | *Terminalia paniculata* | 1158 | 2457 | 0 |
| 6 | Fabaceae | *Gliricidia sepium* | 0 | 28 | 0 |
| 7 | Fabaceae | *Moullava spicata* | 6 | 0 | 0 |
| 8 | Fabaceae | *Xylia xylocarpa* | 19 | 0 | 0 |
| 9 | Loranthaceae | *Dendrophthoe falcata* | 0 | 2168 | 0 |
| 10 | Malvaceae | *Grewia tiliifolia* | 0 | 300 | 0 |
| 11 | Malvaceae | *Helicteres isora* | 18 | 0 | 7 |
| 12 | Rhamnaceae | *Ziziphus rugosa* | 27 | 0 | 0 |
| 13 | Rubiaceae | *Catunaregam spinosa* | 2 | 0 | 0 |
| 14 | Rubiaceae | *Ixora brachiata* | 122 | 25 | 0 |
| 15 | Rubiaceae | *Ixora coccinea* | 0 | 253 | 103 |
| 16 | Rutaceae | *Glycosmis pentaphylla* | 12 | 9 | 0 |
| 17 | Sapotaceae | *Xantolis tomentosa* | 60 | 0 | 0 |
| 18 | Vitaceae | *Leea indica* | 1 | 0 | 0 |

**Table S3.**
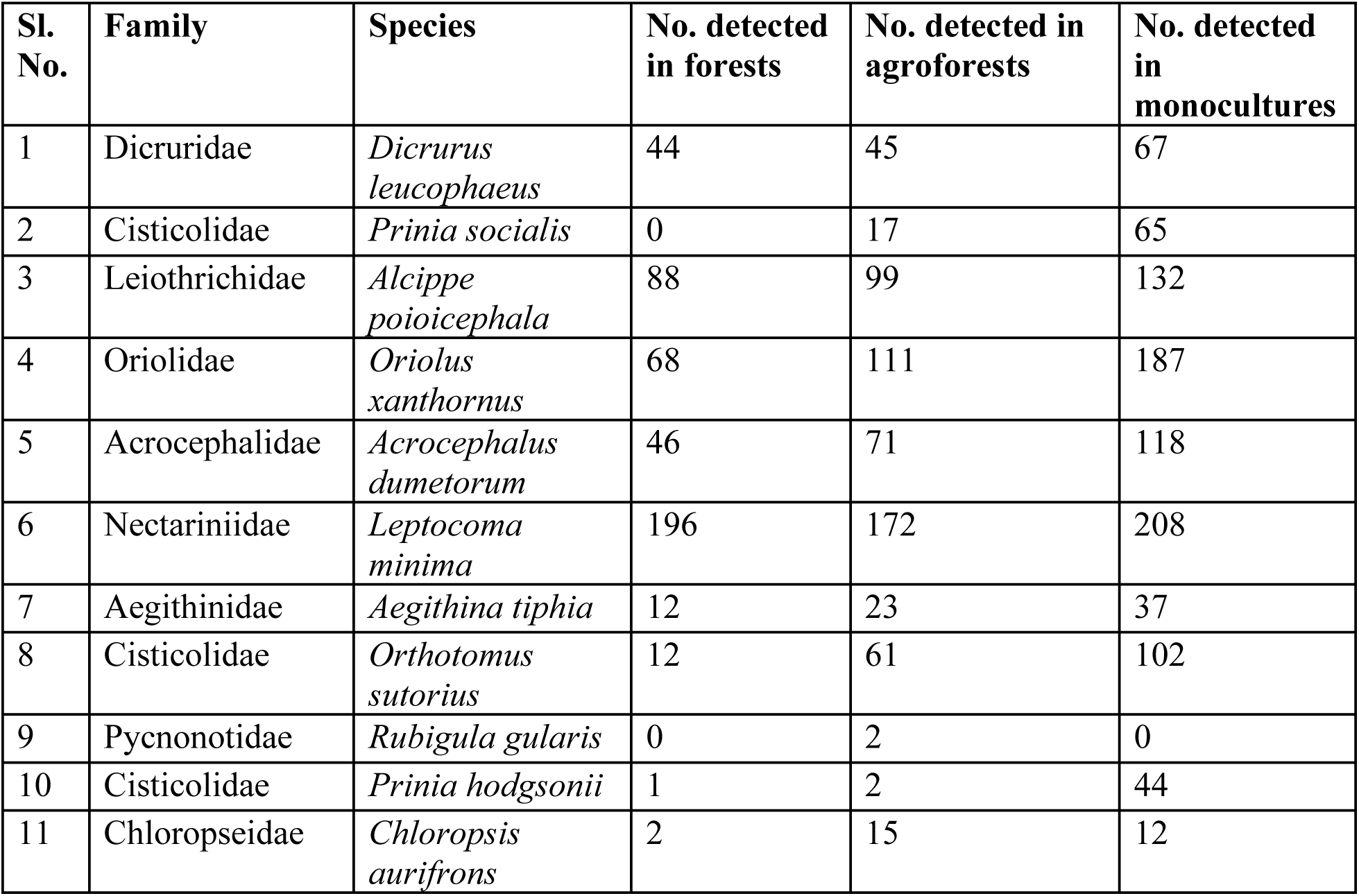
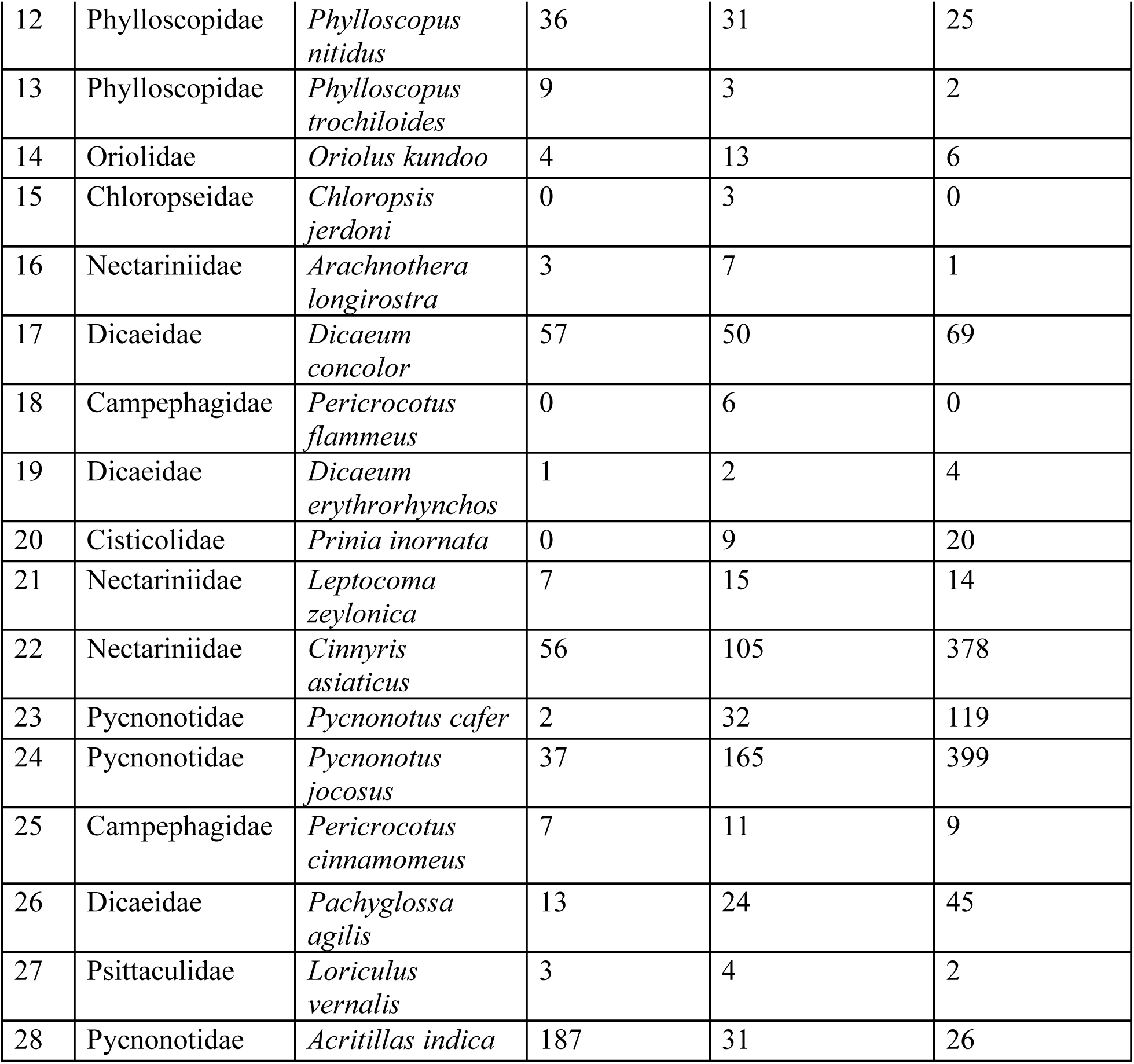
Checklist of bird species that fed on nectar during the sampling period and their abundance across 25 forest points, 25 cashew agroforest points, and 50 cashew monoculture points. 28 of the 31 interacting species were detected in point counts. Species are sorted according to families.

**Table S4.** Checklist of interacting bird species with the number of interactions they participated in, and number of plant species they interacted with. Species are sorted in the decreasing order of the number of interactions.

| Sl. No. | Bird species | No. of interactions | No. of plant species it interacted with |
| --- | --- | --- | --- |
| 1 | *Leptocoma minima* | 274 | 29 |
| 2 | *Cinnyris asiaticus* | 122 | 18 |
| 3 | *Alcippe poioicephala* | 44 | 11 |
| 4 | *Dicaeum concolor* | 37 | 10 |
| 5 | *Leptocoma zeylonica* | 36 | 11 |
| 6 | *Pachyglossa agilis* | 14 | 5 |
| 7 | *Prinia inornata* | 12 | 3 |
| 8 | *Chloropsis aurifrons* | 10 | 5 |
| 9 | *Arachnothera longirostra* | 10 | 3 |
| 10 | *Prinia socialis* | 8 | 4 |
| 11 | *Aethopyga vigorsii* | 8 | 3 |
| 12 | *Prinia hodgsonii* | 6 | 3 |
| 13 | *Oriolus xanthornus* | 5 | 3 |
| 14 | *Pycnonotus jocosus* | 5 | 2 |
| 15 | *Acrocephalus dumetorum* | 4 | 4 |
| 16 | *Orthotomus sutorius* | 4 | 3 |
| 17 | *Rubigula gularis* | 4 | 1 |
| 18 | *Dicrurus leucophaeus* | 3 | 2 |
| 19 | *Chloropsis jerdoni* | 3 | 2 |
| 20 | *Dicaeum erythrorhynchos* | 3 | 2 |
| 21 | *Pericrocotus cinnamomeus* | 3 | 2 |
| 22 | *Loriculus vernalis* | 3 | 2 |
| 23 | *Aegithina tiphia* | 2 | 2 |
| 24 | *Phylloscopus nitidus* | 2 | 1 |
| 25 | *Sturnia blythii* | 2 | 1 |
| 26 | *Acritillas indica* | 2 | 2 |
| 27 | *Phylloscopus trochiloides* | 1 | 1 |
| 28 | *Oriolus kundoo* | 1 | 1 |
| 29 | *Pericrocotus flammeus* | 1 | 1 |
| 30 | *Pycnonotus cafer* | 1 | 1 |

**Table S5.** The effect of time (Julian day of the year) and land-use type on flower abundance from the results of a generalised linear mixed model with zero-inflated negative binomial distribution. Asterisks beside *p*-values indicate estimates whose 95% CIs do not overlap zero and therefore significantly influence flower abundance. *R^2^conditional* = 0.7

| Group | Variance | Standard Deviation | 95% CI |
| --- | --- | --- | --- |
| Point ID | 0.13 | 0.36 | 0.22 – 0.58 |

**Fixed effects**
| Term | Estimate | 95% CI | *p* |
| --- | --- | --- | --- |
| Intercept: Forest | 3.97 | 3.59- 4.35 | < 0.001* |
| Julian day | -0.22 | -0.54 - 0.11 | 0.19 |
| Agroforest | 0.22 | -0.24 - 0.68 | 0.34 |
| Monoculture | -0.31 | -0.73 - 0.1 | 0.14 |
| Julian day : Agroforest | -0.82 | -1.25 - -0.41 | < 0.001* |
| Julian day : Monoculture | -1.57 | -1.94 - -1.21 | < 0.001* |

**Zero-inflation model**
| Term | Estimate | 95% CI | *p* |
| --- | --- | --- | --- |
| Intercept: Forest | 0.88 | 0.5 - 1.27 | < 0.001* |
| Agroforest | -2.51 | -3.17 - -1.84 | < 0.001* |
| Monoculture | -3.55 | -4.73 - -2.38 | < 0.001* |

**Table S6.**
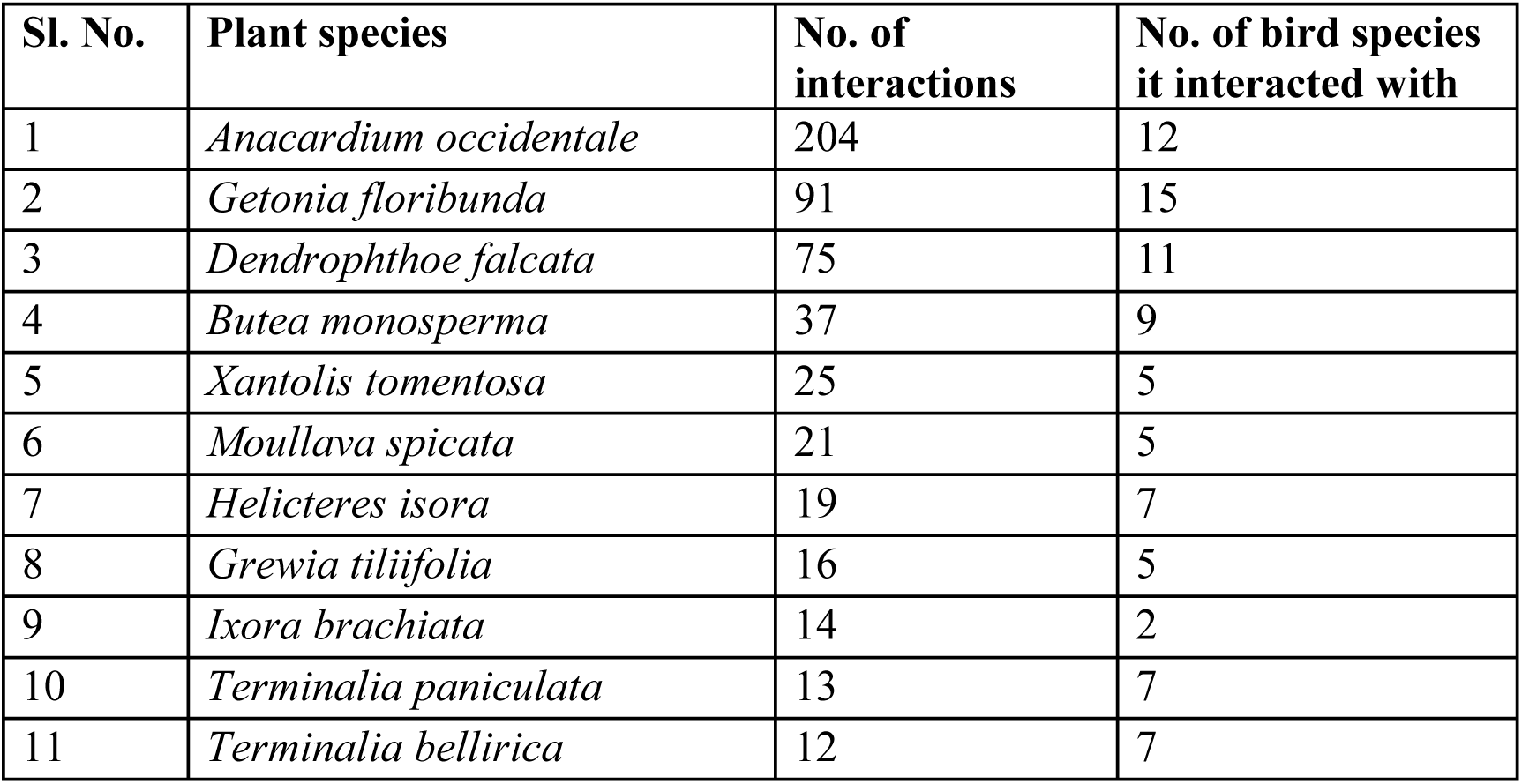
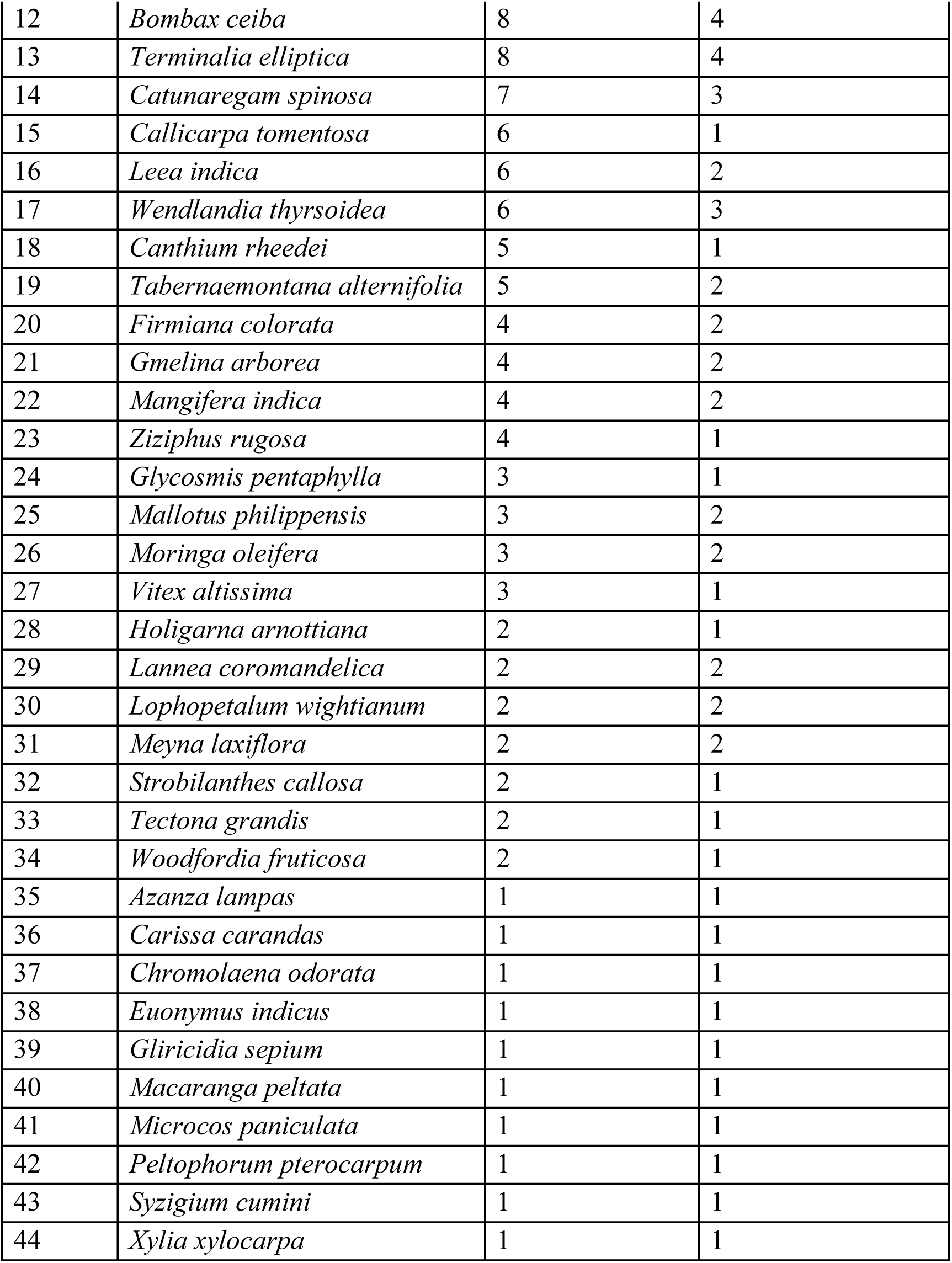
Checklist of interacting plant species with the number of interactions they participated in, and number of bird species they interacted with. Species are sorted in the decreasing order of the number of interactions.

**Table S7.** Multi-model inference table and goodness of fit test results for the distance sampling models used to estimate densities of nectar-feeding birds in forests, agroforests, and monocultures in northern Western Ghats, India. The scale function was modelled as a function of intercept and land-use type (Region.Label).

| Sl. No. | Key function | Formula | $p$ value ($\chi^2$ GoF) | $\hat{P}_a$ | SE ($\hat{P}_a$) | $\Delta$ AIC |
| --- | --- | --- | --- | --- | --- | --- |
| 1 | Half-normal | ∼Region.Label | 0.12 | 0.29 | 0.01 | 0 |
| 2 | Hazard-rate | ∼Region.Label | NA | 0.31 | 0.02 | 6.07 |
| 3 | Half-normal | ∼1 | 0.43 | 0.29 | 0.01 | 9.14 |
| 4 | Hazard-rate | ∼1 | 0.06 | 0.3 | 0.02 | 14.08 |

**Table S8.** Summary of sampling effort and estimated flock densities of nectar-feeding birds in forests, agroforests, and monocultures in northern Western Ghats, India.

| Sl. No. | Land-use type | Effort | Density (flocks/km<sup>2</sup>) | SE | CV | Lower 95% CI | Upper 95% CI |
| --- | --- | --- | --- | --- | --- | --- | --- |
| 1 | Forest | 125 | 9.18 | 0.88 | 0.1 | 7.59 | 11.08 |
| 2 | Agroforest | 125 | 14.99 | 1.38 | 0.09 | 12.49 | 17.98 |
| 3 | Monoculture | 250 | 9.16 | 0.77 | 0.08 | 7.75 | 10.82 |

**Table S9.**
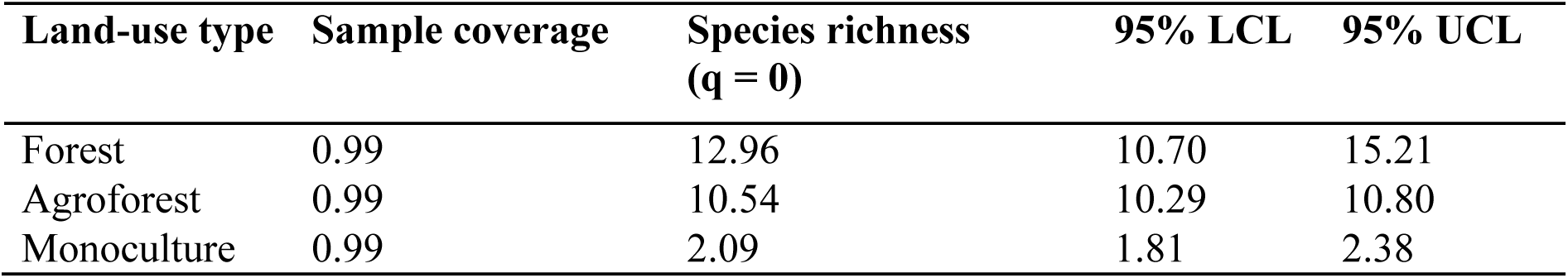
Summary of the sample coverage, species richness values, and associated bootstrapped 95% CI of flowers for the three land-use types (forest, agroforest, and monoculture).

**Table S10.** Summary of the sample coverage, Hill-Shannon (q = 1) numbers, and associated bootstrapped 95% CI of flowers for the three land-use types (forest, agroforest, and monoculture).

| Land-use type | Sample coverage | Hill-Shannon (q = 1) | 95% LCL | 95% UCL |
| --- | --- | --- | --- | --- |
| Forest | 0.99 | 3.55 | 3.38 | 3.71 |
| Agroforest | 0.99 | 3.19 | 3.14 | 3.25 |
| Monoculture | 0.99 | 1 | 0.99 | 1.00 |

**Table S11.** Summary of the sample coverage, species richness (q = 0) values, and associated bootstrapped 95% CI of nectar-feeding birds for the three land-use type (forest, agroforest, and monoculture).

| Land-use type | Sample coverage | Species richness<br>($q = 0$) | 95% LCL | 95% UCL |
| --- | --- | --- | --- | --- |
| Forest | 0.99 | 20.26 | 18.36 | 22.16 |
| Agroforest | 0.99 | 25.39 | 24.02 | 26.75 |
| Monoculture | 0.99 | 20.9 | 19.94 | 21.89 |

**Table S12.** Summary of the sample coverage, Hill-Shannon (q = 1) numbers, and associated bootstrapped 95% CI of nectar-feeding birds for the three land-use types (forest, agroforest, and monoculture).

| Land-use type | Sample coverage | Hill-Shannon<br>($q = 1$) | 95% LCL | 95% UCL |
| --- | --- | --- | --- | --- |
| Forest | 0.99 | 10.82 | 10.14 | 11.5 |
| Agroforest | 0.99 | 15.11 | 14.32 | 15.9 |
| Monoculture | 0.99 | 12.54 | 12.03 | 13.05 |

**Table S13.** Summary of the sample coverage, species richness (q = 0) values, and associated bootstrapped 95% CI of nectarivorous interactions for the three land-use types (forest, agroforest, and monoculture).

| Land-use type | Sample coverage | Species richness<br>($q = 0$) | 95% LCL | 95% UCL |
| --- | --- | --- | --- | --- |
| Forest | 0.71 | 60.63 | 38.21 | 83.05 |
| Agroforest | 0.71 | 38.06 | 26.75 | 49.38 |
| Monoculture | 0.71 | 41.99 | 31.02 | 52.94 |

**Table S14.**
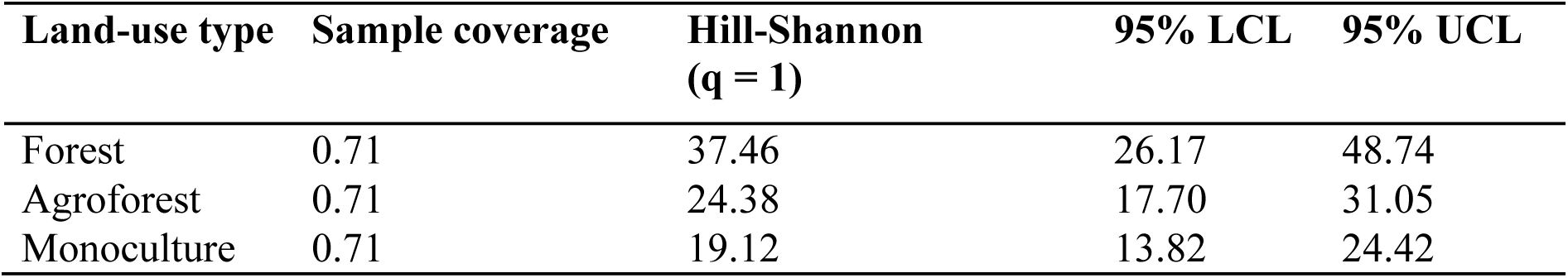
Summary of the sample coverage, Hill-Shannon (q = 1) numbers, and associated bootstrapped 95% CI of nectarivorous interactions for the three land-use types (forest, agroforest, and monoculture).

| Land-use type | Sample coverage | Hill-Shannon<br>($q = 1$) | 95% LCL | 95% UCL |
| --- | --- | --- | --- | --- |
| Forest | 0.71 | 37.46 | 26.17 | 48.74 |
| Agroforest | 0.71 | 24.38 | 17.70 | 31.05 |
| Monoculture | 0.71 | 19.12 | 13.82 | 24.42 |

**Table S15.** Network parameters (Mean ± SD) across land-use types (forest, agroforest, monoculture), after rarefying networks 100 times.

| Network parameter | Forest (Mean ± SD) | Agroforest (Mean ± SD) | Monoculture (Mean ± SD) |
| --- | --- | --- | --- |
| Connectance | 0.12 | $0.14 \pm 0.01$ | $0.163 \pm 0.02$ |
| wNODF | 22.08 | $19.69 \pm 1.89$ | $16.29 \pm 2.56$ |
| $H_2'$ | 0.29 | $0.39 \pm 0.02$ | $0.39 \pm 0.03$ |
| modularity ($Q$) | 0.3 | $0.41 \pm 0.02$ | $0.36 \pm 0.02$ |

